# Precipitation seasonality and soil carbon shape Arcellinida diversity across Bulgaria’s ecosystems

**DOI:** 10.64898/2026.09.29.755277

**Authors:** Ángel García-Bodelón, Milcho Todorov, Nura ElKhouri-Vidarte, Carmen Soler-Zamora, Bertrand Fournier, Enrique Lara

## Abstract

Recent years have witnessed the discovery of an immense diversity of protists in almost all ecosystems on Earth. However, a major challenge consists in identifying diversity patterns and the drivers that explain the distribution of this diversity. While several works have focused on local parameters, only a few studies have addressed this question at regional scales. In this study, we characterize the diversity and distribution of Arcellinida (testate amoebae) at regional scale of a climatically contrasted country, Bulgaria. We collected 75 samples geographically evenly distributed and representative of the country’s natural ecosystems and used metabarcoding to determine their diversity. Communities composition differed markedly among habitats, revealing sharp beta diversity patterns. “Mesophilic” environments such as forests and ponds were most diverse, while acidic and oligotrophic *Sphagnum* peatlands were poorest and hosted the highest proportion of unique taxa. Our analyses of richness-environment based on a Random Forest approach applied to all soil systems indicated that stable humidity (low silt content, low precipitation seasonality) and intermediate soil carbon stocks had a positive effect on richness. The predicted increase in continentality in the next decades in Bulgaria calls for testing climatic models to evaluate the potential risk of declining Arcellinida biodiversity.

## INTRODUCTION

Protists comprise the majority of eukaryotic diversity and play fundamental roles in ecosystem functioning, particularly in soil and freshwater environments where they regulate microbial communities and drive nutrient cycling (Gao et al., 2019; Geisen et al., 2018). Through their interactions with bacteria, fungi, plants and animals, they contribute to key ecosystem services including soil fertility (Xiong et al., 2018) and carbon sequestration (Liao et al., 2024), making them indispensable actors for ecosystem health and human welfare (Berlinches de Gea et al., 2025; Geisen et al., 2018) However, despite their ecological importance, the distribution of their diversity remains mostly unexplored, especially at regional scales (i.e. country-wide) (Geisen et al., 2017).

One of the major challenges lies in the immense diversity and ecological complexity of protists. Closely related taxa can differ markedly in their trophic strategies (Glücksman et al., 2010) and environmental preferences (Singer et al., 2018), and a large fraction of their diversity remains undescribed. Traditional morphology-based approaches have substantially underestimated this diversity due to cryptic species and convergent traits (González-Miguéns et al., 2022; Soler-Zamora et al., 2023), while recent advances in environmental DNA (eDNA) sequencing have revealed a much finer-scale structuring of protist communities (Pawlowski et al., 2012). Diversity surveys based on eDNA allow a fast and accurate assessment of protist diversity in large sampling designs, and open the way to geographical upscaling. Yet, even with these tools, linking community composition to environmental drivers in a predictive framework still remains difficult. Such knowledge would allow foreseeing the response of protist communities to ongoing climate change, which remains poorly understood, limiting our ability to predict how ecosystem processes will shift in the coming decades.

Ongoing research is nevertheless providing promising results. Regional scale, eDNA-based studies revealed the prevalence of topoclimatic factors on protist diversity distribution in the Swiss Alps (Seppey et al., 2020) or at global scale (Oliverio et al., 2020). Climatic parameters such as temperature and humidity (Fernández et al., 2022) or landscape structure (Seppey et al., 2023) have been found to drive diversity at a regional scale. However, the great differences in size, lifestyle, ecological function, and even genetic variability among protists make it considerably difficult to draw any general conclusion (Lara et al., 2022). Indeed, recent studies show that, in soil, community assembly rules change according to the organisms functions (Fernández et al., 2022).

Focusing on ecologically coherent model groups offers a promising way forward. Arcellinida (Amoebozoa) are particularly suitable in this regard. As predominantly eukaryvorous protists, they occupy high trophic positions within microbial food webs and are tightly linked to the composition of surrounding microeukaryotic communities (Song et al., 2018). Many species exhibit narrow ecological tolerances with respect to moisture, temperature, and nutrient availability, and their test morphology reflects adaptations to environmental constraints such as desiccation (Fournier et al., 2012). As a result, Arcellinida community composition is highly responsive to environmental gradients (ElKhouri-Vidarte et al., 2026), making them effective indicators of ecosystem conditions. Recent metabarcoding approaches now allow their diversity to be assessed with high resolution across a wide range of habitats (González-Miguéns et al., 2023), making regional scale studies possible.

In this study, we investigated Arcellinida diversity across a broad range of ecosystems at the scale of an entire country, Bulgaria, encompassing strong climatic and environmental gradients. Specifically, our objectives were to (i) characterize patterns of alpha and beta diversity across habitat types, hypothesizing that species richness would differ among habitats and that community composition would cluster according to habitat type; and (ii) identify the main climatic and edaphic drivers of community richness. Based on these findings, we suggest a possible influence of climate change on future species richness, following the predictions made for Bulgaria. This influence is attributed to an increase in precipitation seasonality caused by the northward extension of the Mediterranean climate zone. Therefore, our study aims to provide a baseline for anticipating how protist communities, and the ecological processes they support, may respond to future climate change.

## MATERIAL AND METHODS

### Study area and sampling

For this study, during the period from July to November 2022, we collected 75 samples across the entire territory of Bulgaria, representing the country’s main natural ecosystems in a geographically balanced way. The sampled habitats included stagnant freshwater bodies, wet and *Sphagnum* mosses, soils, litter and mosses in deciduous and coniferous forest, soils and mosses in xerothermic habitats and sandy interstitial on the seashore (Fig. 1). In this way, we aimed to identify differences in diversity and species richness between different habitats. Sample collection in different environments differs and was carried out as follows:

**Figure 1.**
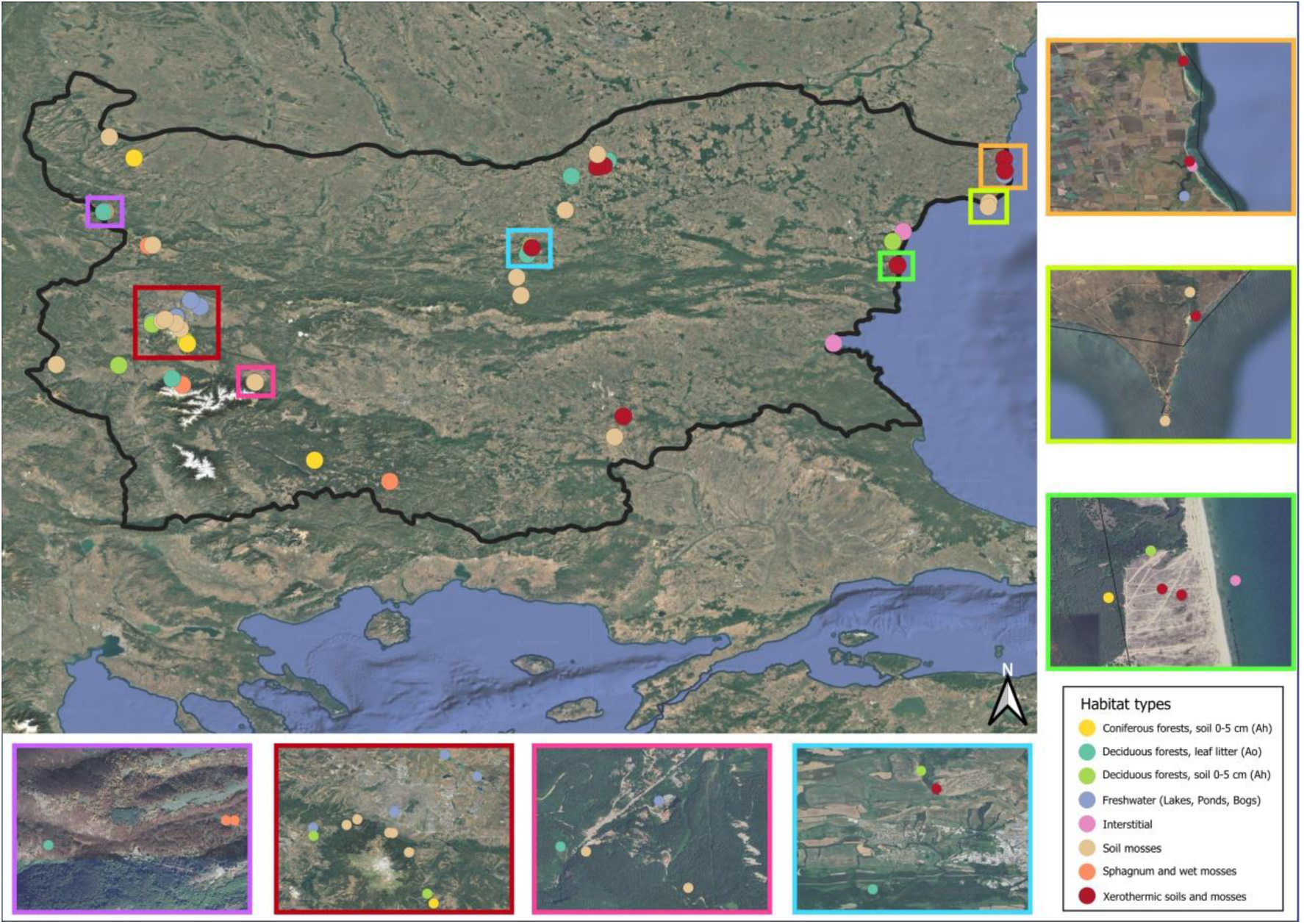
Map of sampling locations in Bulgaria, identified by colour circles. Colours indicate ecosystem types. The framed insets act as magnified views (zoomed areas) to show certain sampling sites in greater detail, using a colour scheme.

• Aquatic samples were obtained by washing aquatic plants in a pail with water from the reservoir itself. Benthic samples were collected using a bottom dredge. The collected sample was allowed to settle for 3-5 minutes, after which the supernatant was removed and the precipitate was filtered through a 500 µm mesh sieve. After sedimentation of the filtrate, 2 samples of about 25-30 ml were taken in Falcon tubes, and fixed into Qiagen’s Lifeguard solution and used for molecular work.

• Terrestrial samples consisted in 400-500 g of fresh material (litter/moss) washed and shaken into 1 l of tap water, filtered through a 500 µm mesh sieve and allowed to settle for 30 minutes. After sedimentation of the filtrate, 2 samples of about 25-30 ml were taken, one of which was fixed with 2% formalin for observation and the other for molecular work.

• The interstitial samples were collected from the Black Sea beaches, by digging holes in the sand about 3-5 m from the waterline, with a depth of 50-80 cm and taking a 500 ml sample from the bottom of the hole in a ratio of 1/2 groundwater and 1/2 sand. Samples from this environment were not fixed and were examined live in laboratory conditions before being stored for further DNA work.

Sampling completeness was calculated as the ratio between observed species richness and bootstrap-estimated richness using the specpool() function from the R package vegan (Oksanen et al., 2026), which estimates total species richness based on bootstrap resampling. Here, sampling completeness was high, with a sampling completeness of 83.7% (Fig. S1, S2).

### eDNA extraction, PCR and sequencing

Total eDNA was extracted using the Qiagen DNeasy PowerSoil Pro Kit. We employed a two-step nested PCR protocol specifically designed for Arcellinida to amplify the cytochrome oxidase subunit I (COI) region, as described by González-Miguéns et al. (2023). We performed the first PCR with universal COI primers (Folmer et al., 1994), and the second PCR with a specific Arcellinida primer (González-Miguéns et al., 2022a) with an unique combination of tags for each sample to identify them. Amplified products were quantified using an Invitrogen Qubit 3 fluorometer with a Thermo Fisher Scientific dsDNA High Sensitivity assay kit. Finally, purified DNA products were normalized, pooled and sequenced on an Illumina NextSeq 2000 platform using 600 cycles at the Genomic Unit of the Fundación Parque Científico de Madrid (Spain).

### eDNA Data curation and “tag jumping”

Illumina reads were curated following this pipeline based on the previous work of González-Miguéns et al. (2023): (1) trimming of primers and demultiplexing was performed using Cutadapt ver. 2.8 (Martin, 2011) following the pipeline “Cutadapt_Pipeline.bash”. (2) The resulting reads per sample were analyzed with the DADA2 R package (Callahan et al., 2016), filtering and trimming after positions 210 and 200 for the forward and reverse respectively. We dereplicated them, merged the paired reads, generating amplicon sequencing variants (ASVs), and removed the chimeric sequences. (3) For ASVs taxonomic assignation we applied VSEARCH ver. 2.14 (Rognes et al., 2016), using the eKOI database (González-Miguéns et al., 2025).

Samples were bioinformatically sorted using sample-specific indexes (“tags”), which implies a risk of sample misassignment (“tag jumping”) (Rodriguez-Martinez et al., 2023; Schnell et al., 2015). To reduce this bias, we carried out a combination of protocols developed in earlier studies (ElKhouri-Vidarte et al., 2026; González-Miguéns et al., 2023; Useros et al., 2024). The necessary thresholds to remove misassigned sequences were determined following the protocol described in (ElKhouri-Vidarte et al., 2026). Briefly, this method was based on the observation of tag combinations that do not correspond to any sample, and are therefore necessarily produced by tag jumping. The proportion of these “false reads” was used to set a threshold. In a first step, we eliminated all ASVs below a minimum read count of 15 thus excluding those with too few reads that might be associated with errors or noise. Subsequently, we calculated the thresholds based on read proportions between unused tag combinations and “real” samples. Thereafter, data were filtered to retain only ASVs assigned to the order Arcellinida with a percentage identity ≥84%. This number was determined empirically in previous studies (González-Miguéns et al., 2024).

### ASVs clustering into OTUs

The ASVs assigned to Arcellinida were aligned with the Arcellinida COI sequences data of González-Miguéns et al. (2022b) and the metabarcoding data of González-Miguéns et al. (2024) using the MAFFT auto algorithm (Katoh et al., 2002) as implemented in Geneious version 2019.0.4.

In order to approximate as much as possible our diversity inventory to species composition, we grouped ASVs into operational taxonomic units (OTUs) using a distance matrix obtained with R package DECIPHER ver. 2.22 (Wright, 2016). We set a barcoding gap of 3% divergence as defined in earlier works (ElKhouri-Vidarte et al., 2026; García-Bodelón et al., 2024). This barcoding gap corresponds to the distance between the sister species *Arcella salobris* and *Arcella uspiensis*, which differ deeply in their tolerance to salt (Useros et al., 2023). The ASVs were clustered into OTUs using the R package DECIPHER ver. 2.22 (Wright, 2016), using the “complete method”, meaning that the maximum distance between any pair of ASVs from a cluster is 0.03.

### OTU richness and community composition

The number of OTUs per sample was use as a proxy for species richness. This metric is widely used in metabarcoding studies as a direct and interpretable measure of diversity (ElKhouri-Vidarte et al., 2026; García-Bodelón et al., 2024; Mahé et al., 2017; Shin et al., 2016; Weydmann-Zwolicka et al., 2024). To ensure the validity of this proxy, we tested the relationship between OTU richness and sequencing depth using Pearson’s correlation coefficient and found no significant linear correlation (r = 0.11, p = 0.37).

In order to compare the observed richness differences across habitat types, we performed Kruskal-Wallis tests, followed by a Dunn test through the FSA package (Ogle et al., 2025) with a Benjamini-Hochberg correction between each pair of habitat types. We performed a non-metric multidimensional scaling (NMDS) to illustrate beta diversity patterns among communities associated to each sampling site. The analysis was based on presence–absence Jaccard distances and illustrated using also ggplot2 using a pairwise 2D projections of the NMDS axes (k = 3) to visualise clearly sample clustering across a three-dimensional ordination than the classical bi-dimensional representation. All figures were edited using Adobe Illustrator CS4 software ver. 14.0.0 (Adobe, 2008).

### Environmental data

We obtained 34 site-specific bioclimatic variables of ∼1 km resolution raster from CHELSA v2.1 (Brun et al., 2022; Karger et al., 2017) and 11 site-specific edaphic variables from ISRIC database https://files.isric.org/soilgrids/latest/data of SoilGrids 2.0 (Poggio et al., 2021) with a resolution of 1 km to capture the influence of terrestrial substrate on microbial habitats (Table S3). To reduce multicollinearity and avoid including highly redundant predictors, highly correlated variables were removed following Dormann et al. (2013), using pairwise Spearman correlations with |r| > 0.7. A correlation matrix was calculated using only pairwise complete observations to ensure that each correlation coefficient was based on all available data for each variable pair. Highly correlated variables were filtered using the findCorrelation() function from the caret R package (Kuhn, 2008), which iteratively removes variables involved in high pairwise correlations by excluding those with the highest mean absolute correlation across all predictors, thereby retaining a subset of minimally redundant variables (Table 1).

**Table 1.** Environmental variables retained after correlation analysis.

| Variable names | Abbreviations |
| --- | --- |
| Temperature of the growing season | gst |
| Mean temperature of the driest quarter | bio9 |
| Precipitation seasonality | bio15 |
| Surface downwelling shortwave flux in air | rsds |
| Net primary production | npp |
| Silt content | silt |
| Coarse fragments volumetric fraction | cfvo |
| Cation exchange capacity buffered at pH 7 | cec |
| Organic carbon stocks | ocs |
| Organic carbon density | ocd |

### Linking Community Composition (beta diversity) to Environmental Drivers

The relationship between microbial community structure (NMDS ordination of the presence–absence matrix with Jaccard distance) and the filtered environmental variables was assessed using the envfit() function from the vegan R package (Oksanen et al., 2026). Significance of environmental vectors was evaluated using permutation tests (999 permutations), and significant vectors (p < 0.05) were scaled and overlaid on the NMDS plot to visualize their influence on community composition.

### Random forest and partial dependence analysis

We assessed the influence of environmental predictors on species richness (as number of OTUs) for all habitat types except freshwater and interstitial habitats, using Random Forest models fitted with the randomForest package (Breiman, 2001). The model was built using only the 60 terrestrial samples, using the previously filtered set of environmental variables (Table 1). To identify the most informative predictors while accounting for spatial autocorrelation, we performed forward feature selection (FFS) using the ffs() function from the CAST package (Meyer et al., 2026). During feature selection, Random Forest models were fitted with 500 trees, 2 variables randomly sampled at each split, and a minimum terminal node size of 15. The FFS procedure selected organic carbon stocks (ocs), silt content (silt), and precipitation seasonality (bio15).

The final Random Forest model was fitted using the selected predictors with 500 trees, 2 variables randomly sampled at each split, a minimum terminal node size of 15, and a bootstrap sampling size of 20 observations. Spatial cross-validation was implemented by clustering the sample coordinates into four spatial folds using k-means clustering in UTM coordinates (10 repetitions of the four-fold spatial partitioning). For each repetition, the model were trained on 3 spatial clusters and tested on the remaining cluster. Predictive performance was quantified using R², root mean square error (RMSE), and mean absolute error (MAE). Variable importance was assessed using permutation importance (%IncMSE) computed with the importance() function from the randomForest package.

Partial dependence plots (PDPs) were generated for the most informative combination of predictors, selected by FFS procedure, on the estimated response (OTU richness), while maintaining the other predictors in their observed distribution. Partial dependence was computed using the partial() function from the pdp R package (Greenwell, 2017). The uncertainty of the partial dependence plots was estimated via bootstrap aggregating (n=50). The 2.5th and 97.5th percentiles of the bootstrap predictions define the uncertainty envelope.

All analyses were conducted in R version 4.5.1 using a fixed random seed (set.seed(123)).

## RESULTS

### Arcellinida diversity

High-throughput, Arcellinida-specific sequencing of the mitochondrial COI barcode region yielded 37 million quality-filtered reads. The number of Illumina reads per sample through DADA2 pipeline is detailed in Table S1. The stepwise filtering approach resulted in a high-confidence dataset of 4273 total ASVs clustered into 545 OTUs (Table 2). We retrieved sequences representative of all higher ranking Arcellinida taxa with the exception of Organoconcha.

**Table 2.**
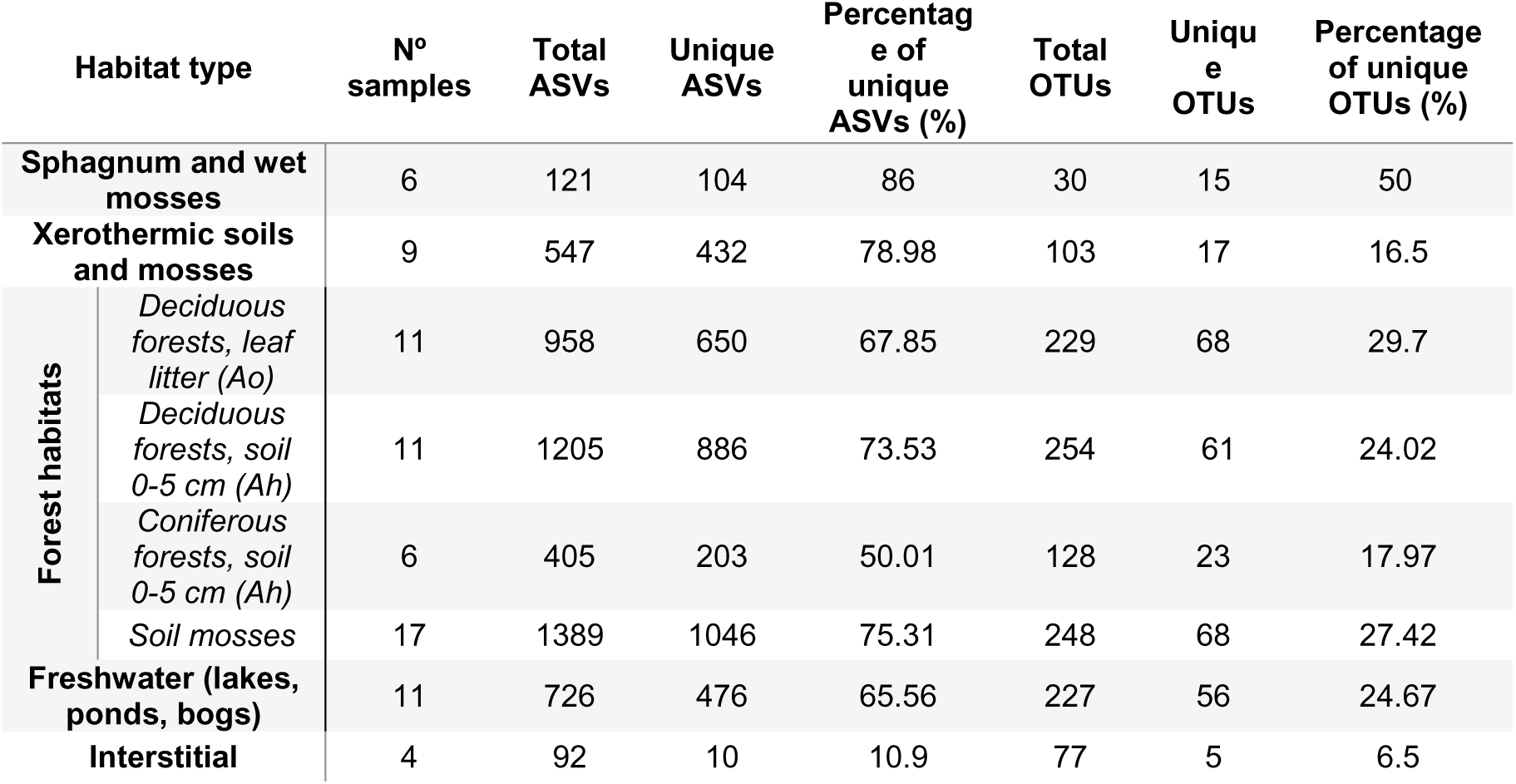
Number of unique ASVs and OTUs, total ASVs and OTUs and percentage of unique ASVs and unique OTUs (%) of each habitat type.

### Alpha diversity and OTU richness

Forest habitats (coniferous, deciduous forests and forest soil mosses) and freshwater exhibited the highest OTU richness, with high variability among samples, with no significant differences between them. In contrast, xerothermic mosses, *Sphagnum* and interstitial habitats were significantly less diverse, being *Sphagnum* the lowest one (Fig. 2).

**Figure 2.**
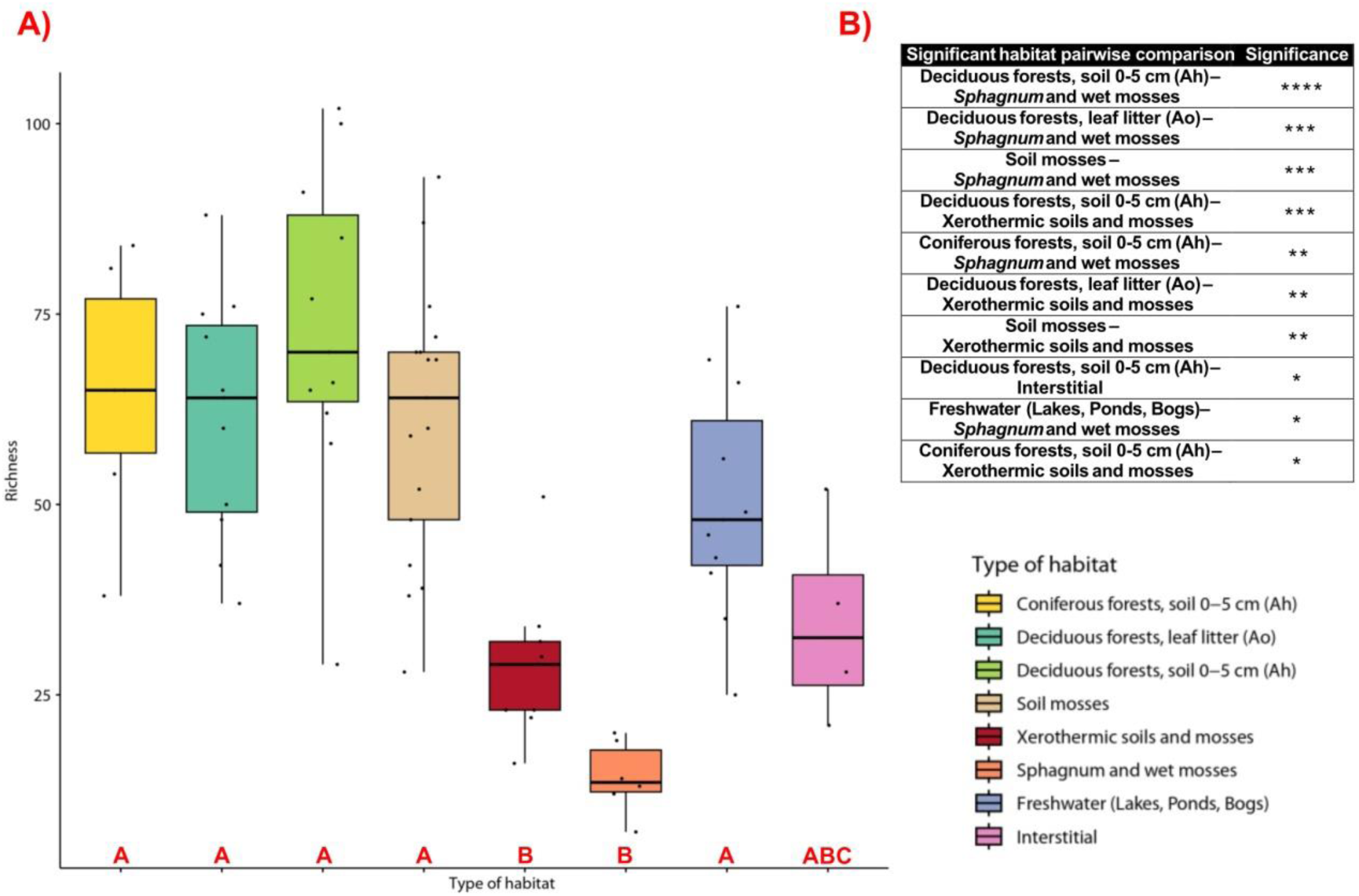
A) Boxplot of species richness (number of total OTUs) across different habitat types. The plot shows the mean species richness calculated from individual samples (dots) within each habitat. The boxes represent the interquartile range of richness, with the central line indicating the median. Error bars represent the interquartile range. Each habitat is represented with a unique colour as shown in the legend. Red letters indicate the significant differences in richness (same letter, no significant difference). B) Significant pairwise comparisons of species richness medians across habitat types, assessed using the Kruskal-Wallis followed by a Dunn test with a Benjamini-Hochberg correction. Significance levels are indicated as follows: **** (p ≤ 0.0001), *** (p ≤ 0.001), ** (p ≤ 0.01), * (p ≤ 0.05).

The uniqueness of associated communities varied among habitat (i.e. unique ASVs and OTUs in each habitat) (Table 2). For ASVs, the highest percentage of unique ASVs was recorded in *Sphagnum* (86.00%), followed by xerothermic soils and mosses (78.98%) and soil mosses (75.31%). In contrast, the lowest percentage was observed in the interstitial habitat (10.90%). Accordingly, the highest percentage of unique OTUs was also found in *Sphagnum* and wet mosses (50.00%), while the interstitial habitat had the lowest percentage (6.50%) as well (Table 2).

The NMDS analysis graphically illustrates community clustering according to their corresponding habitat. While forest habitats formed a single cohesive group, freshwater, xerothermic mosses and *Sphagnum* appeared as unique clusters and did not share statistical space with each other. However, interstitial communities clustered together with forest samples (Fig. 3).

**Figure 3.**
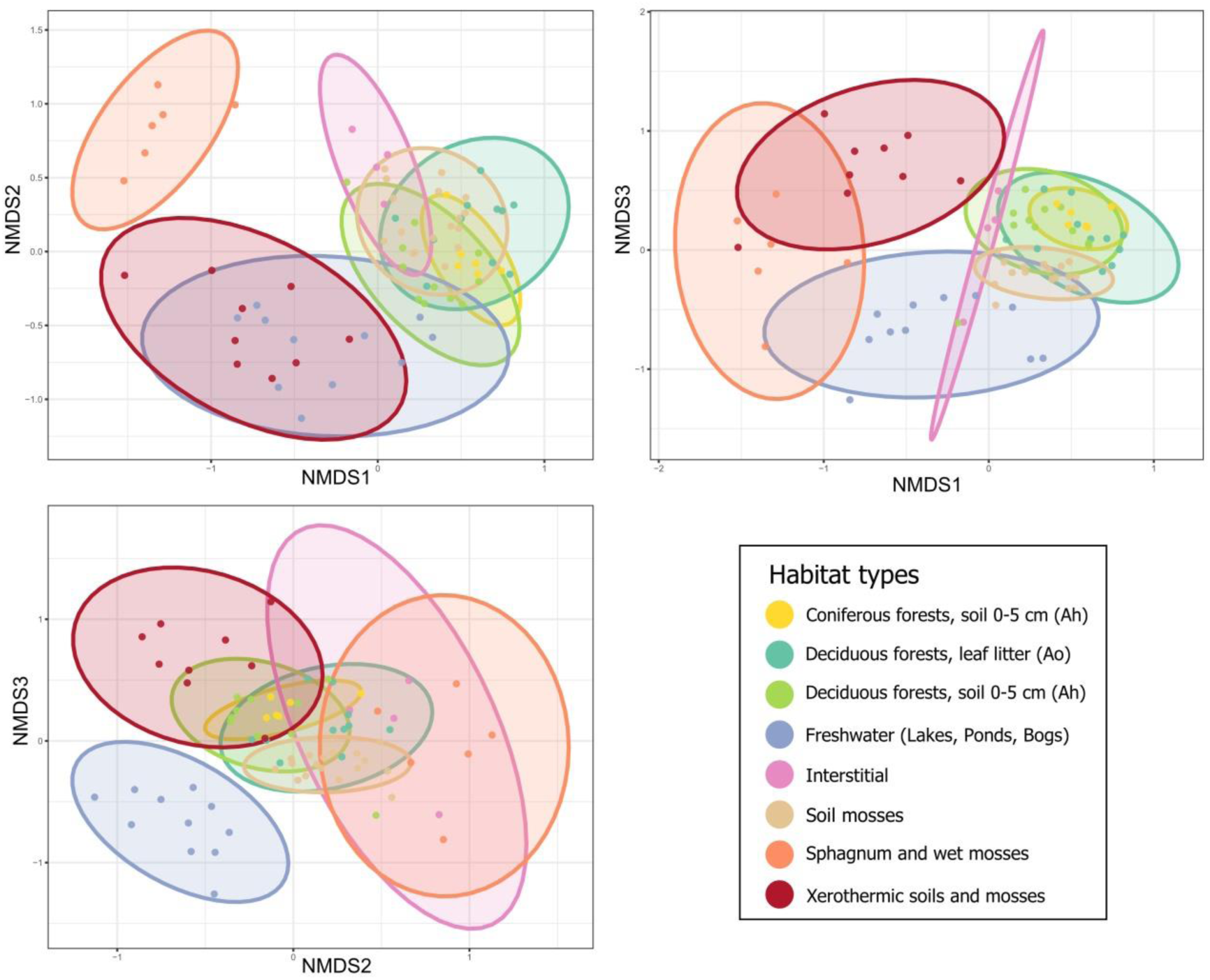
Non-metric multidimensional scaling (NMDS) plot based on Jaccard distance calculated from presence–absence data for the eight habitat types (k = 3, stress = 0.154). Each point represents a sample, and habitats are differentiated by colour according to the legend. The three 2D projections of the NMDS axes (NMDS1 vs NMDS2, NMDS1 vs NMDS3, NMDS2 vs NMDS3) are shown to facilitate interpretation of relationships between samples.

The envfit analysis revealed significant correlations between the NMDS ordination and several environmental variables obtained from the CHELSA and ISRIC databases (Fig. 4).

**Figure 4.**
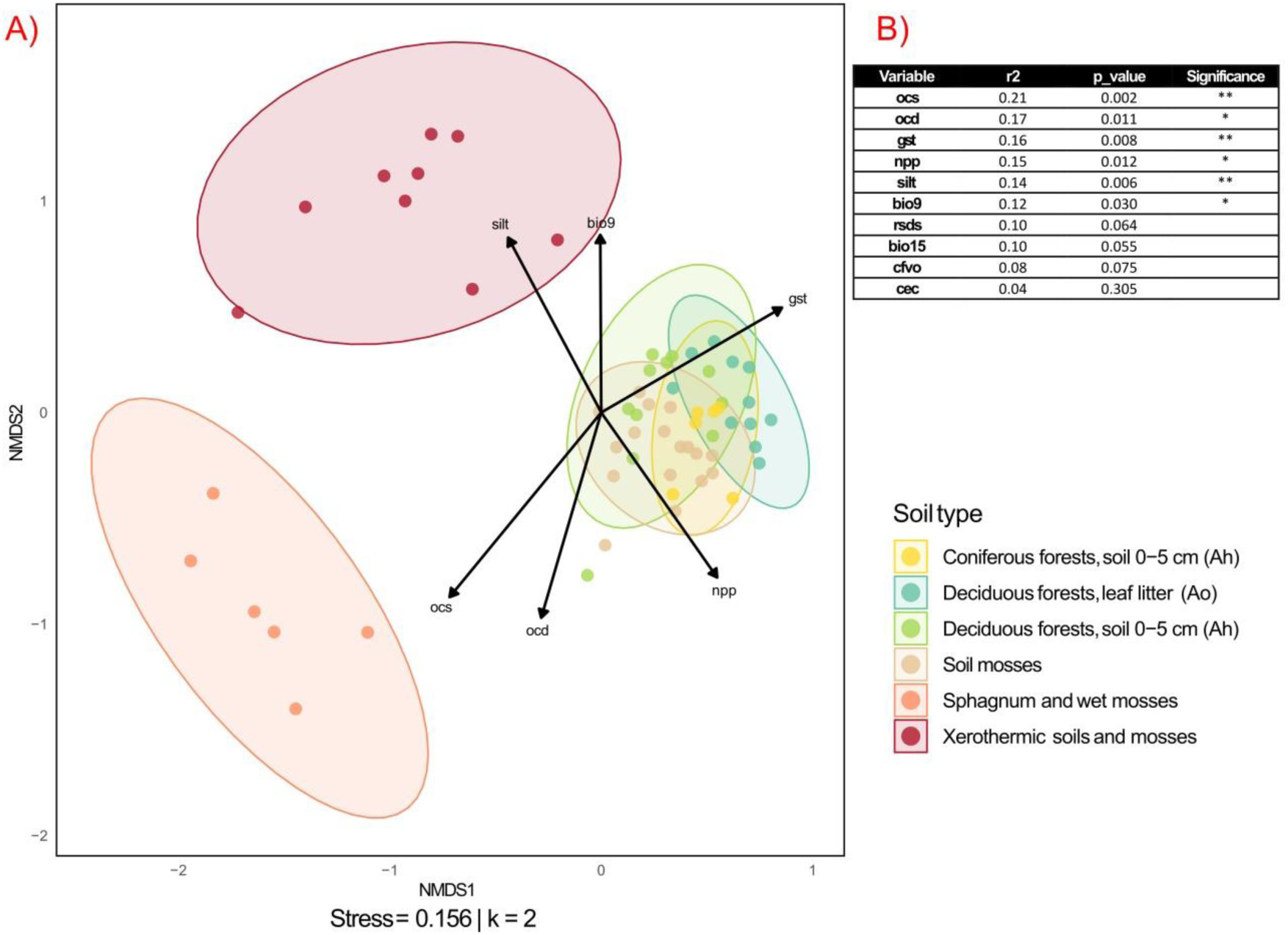
A) Non-metric multidimensional scaling (NMDS) based on a terrestrial community matrix using Jaccard dissimilarity (k = 2, Stress = 0.156). Only soil habitats are shown; freshwater and interstitial habitats were excluded from the analysis. The points represent samples coloured according to habitat type. Confidence ellipses (95%) show the variation within each habitat. The arrows indicate the direction and magnitude of significant environmental variables (p < 0.05). B) Significant environmental variables identified by the envfit analysis (p < 0.05). Significance levels are indicated as follows: ** (p ≤ 0.01), * (p ≤ 0.05).

### Modeling OTU richness

Forward feature selection (FFS) identified 3 environmental predictors as the most informative variables explaining variation in OTU richness: organic carbon stocks (ocs), silt content (silt), and precipitation seasonality (bio15); whereas the addition of further predictors did not improve model performance. The final Random Forest model explained approximately 20% of the spatial variation in OTU richness (Spatial cross-validation R²= 0.20), with a mean RMSE of 24.93 ± 0.14 and a mean MAE of 20.81 ± 0.07 (Table 2).

**Table 2.** Random Forest metrics used: coefficient of determination (R²), root mean square error (RMSE), and mean absolute error (MAE). Predictor importance was assessed using the percentage increase in mean squared error (%IncMSE), with higher values indicating greater contribution to model accuracy. Values are presented as mean ± standard deviation across cross-validation folds.

| Metric | Value |
| --- | --- |
| Spatial cross-validation $R^2$ | $0.20 \pm 0.01$ |
| Train $R^2$ | $0.36 \pm 0.01$ |
| RMSE | $24.93 \pm 0.14$ |
| MAE | $20.81 \pm 0.07$ |
| Predictors (%IncMSE) | ocs (14.71); bio15 (8.18); silt (3.66) |

Partial dependence plots (PDPs) revealed decreasing non-linear responses of richness to the examined environmental gradients (Fig. 5). Organic carbon stocks (ocs) exhibited maximum predicted values at intermediate carbon concentrations (4.5–5.2 kg m⁻²), followed by a pronounced decline reaching a plateau at higher levels (Fig. 5A). Precipitation seasonality (bio15) displayed a predominantly negative relationship with richness, declining progressively with the highest values associated with more climatically stable conditions (Fig. 5B). Finally, richness decreased with silt content (silt), with steeper slopes at the lowest and highest values (Fig. 5C).

**Figure 5.**
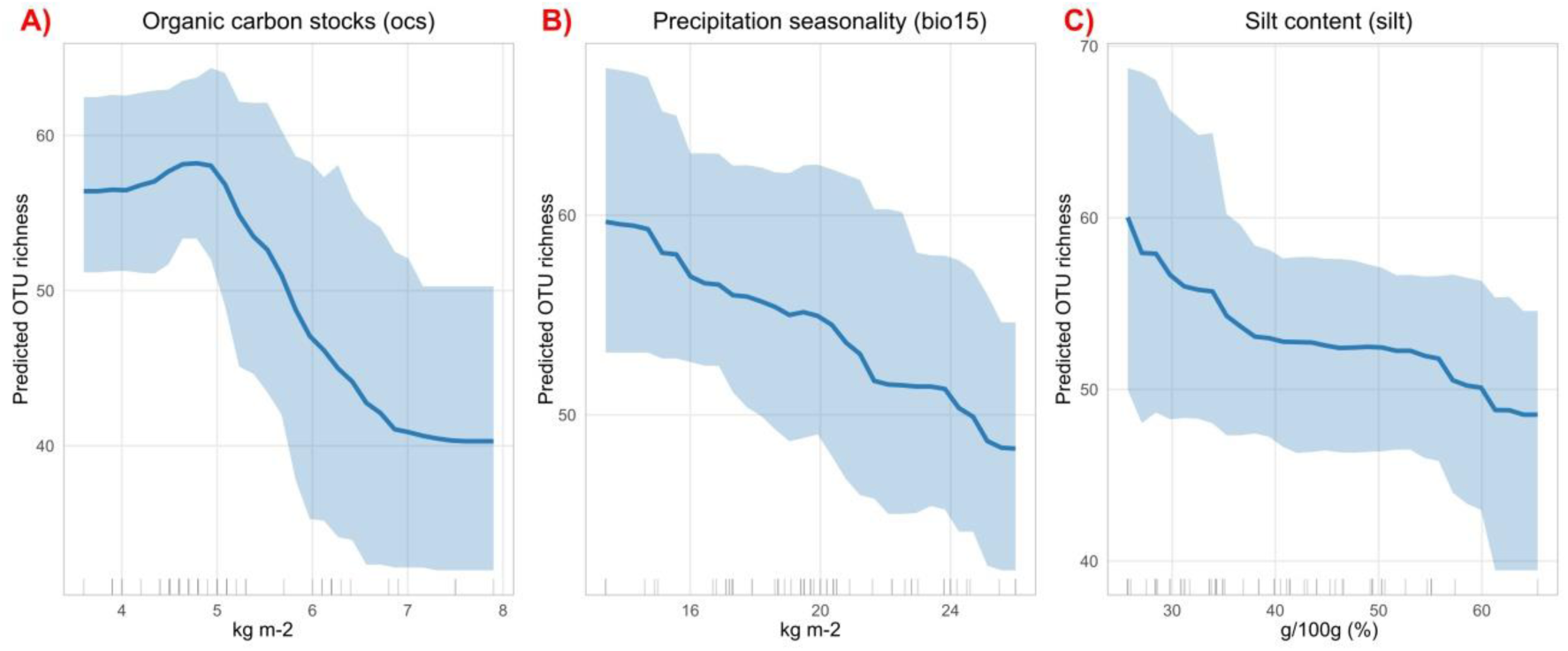
Partial dependence plots (PDPs) from the Random Forest model showing the effects of the predictors selected through the forward feature selection (FFS) procedure: A) organic carbon stocks (ocs), B) precipitation seasonality (bio15), and C) silt content (silt). Shaded areas represent 95% uncertainty bands from 50 bootstrap replicates of the Random Forest model. Rug plots along the x-axis show the distribution of observed values (n = 60) for each predictor.

## DISCUSSION

### Richness patterns in the different habitats

Arcellinids are ecologically restricted in terms of humidity, temperature and nutrient availability (Fournier et al., 2012), in line with the differences detected in richness and OTU composition observed between habitats. Our analyses revealed the highest diversity in forest habitats, as well as in freshwater. In both forest soils and freshwater habitats, richness varied extensively, from ≈25 OTUs to over 100 (Fig. 2). These habitats are relatively mesophilic (in terms of temperature, or pH) thus permitting the development of a high diversity. Forest habitats (mosses and litter) hosted a large amount of Excentrostoma, including many OTUs from genera *Centropyxis* and *Plagiopyxis* (Table S2). Such organisms have been suggested to be either phenotypically very variables, or extremely diverse (Foissner and Korganova, 2000). Our results, as well as other molecular-based works (ElKhouri-Vidarte et al., 2026; González Miguéns et al., 2023), suggests that they are actually hyperdiverse soil inhabitants. Freshwater environments included organisms with a great variety of test outlines, which suggests many different ecological functions (Fournier et al., 2012). Indeed, we detected OTUs from the genera *Cucurbitella*, *Cylindrifflugia*, *Difflugia*, *Galeripora* and *Phryganella* (Table S2), which differ deeply both through their morphology and their lifestyles (Iranzo-Cano et al., 2026). The large number of available microhabitats — such as sediments, aquatic vegetation and plankton — exerts divergent pressures on organisms and promotes functional diversification.

In contrast, *Sphagnum* and xerothermic habitats appeared to host a significantly lower diversity (Fig. 2). In *Sphagnum*, low pH, low nutrient amounts and high proportion of recalcitrant carbon act as important ecological filters on associated organisms (Dedysh et al., 2006). In xerothermic habitats, high temperatures and desiccation might also play a similar role in selecting only those taxa capable of surviving under harsh conditions. In contrast to forest soil habitats and freshwater, richness was relatively homogeneous in both Sphagnum and xerothermic habitats (Fig. 2). The interstitial habitats also showed reduced richness (77 OTUs), albeit more heterogeneous than *Sphagnum* and xerothermic habitats. Such depleted diversity is in line with Useros et al. (2025) who showed that marine influenced habitats hosted almost no Arcellinida.

### Distinctiveness of habitat communities

Patterns observed in NMDS analyses show that while forest soil habitats group together as a single cluster, communities from freshwater, xerothermic mosses and *Sphagnum* cluster separately. Interstitial habitat communities cluster close to forest soil (Fig. 3 and 4). This pattern suggests that forest communities do not differ fundamentally from each other. However, coniferous forests occupy a smaller statistical space than deciduous forests soil in the NMDS, which suggest more homogeneous community composition. In turn, freshwater, xerothermic and *Sphagnum* communities appear very distinct in the NMDS, with no overlap. Freshwater and terrestrial communities differ deeply in their composition, as already shown in earlier metabarcoding-based studies (González-Miguéns et al., 2024). Such results advocate for different selective pressures exerted on communities with respect to forest habitats, and a tendency for organisms to be highly specialized to certain ecosystems.

In order to better characterize the level of specialization of the communities, we compared the number of OTUs that appeared exclusively in a single habitat type. *Sphagnum* and wet mosses exhibited the highest number of habitat-specific organisms, reaching a percentage of 50% of all its OTUs (Table 2). Low diversity, highly differentiated communities and a high number of exclusive taxa suggest that *Sphagnum* mosses create a highly selective environment for Arcellinida, likely driving taxa adaptation and evolution (Singer et al., 2018b). *Nebela collaris*, which appeared in this study (Table S2), are an example of the diversity typically found in these well-described assemblages (Mitchell et al., 2008). Freshwater systems had a lower proportion of unique OTUs (27.4%). The fact that samples were collected near the shore, which represents an ecotone zone where terrestrial and freshwater species tend to mix (González-Miguéns et al., 2024) is probably the reason for the relatively low number of exclusive OTUs. Genus *Difflugia*, detected in this study (Table S2), typically lives in freshwater systems.

In spite of their low diversity and strong environmental pressure, xerothermic soil and mosses had relatively few unique OTUs (16.5%) in contrast to *Sphagnum* habitats. This might suggest that most local species did not manage to adapt to these ecosystems, and communities are mostly formed by those species that can stand desiccation and heat. Nevertheless, an OTU corresponding (99.1%) to the species *Heleopera steppica*, first discovered in Mediterranean-continental shrublands (Ribeiro et al., 2023) can be found there (Table S2). This species has a slit-like aperture, a trait thought to protect the organism against dessication (Bonnet, 1961). Finally, interstitial habitats had the lowest proportion of unique OTUs (6.5%) (Table 2). Its low richness (Figure 2), grouping with forest soil samples and heterogeneity in the NMDS could suggest that most species present derive from other habitats. This conclusion supports the hypothesis in Useros et al., (2025) that Arcellinida never truly managed to colonize the sea, and that organisms that live in marine influenced habitats are derived from continental ecosystems.

### Diversity drivers and global change indicators

To better understand the selective pressures exerted on the communities, we determined the main drivers of beta diversity. Parameters related to soil carbon content (ocs, ocd), climate (gst, bio9), physical soil characteristics (silt), and productivity (npp) had the most important influence on communities composition. Altogether, metrics related to water availability/evaporation and nutrient turnover were the drivers that best explained community composition (Fig. 4); unlike other studies on protist soil diversity, pH did not seem to play a significant role in diversity distribution (Geisen et al., 2015). These results highlight the existence of taxa that are specialized in a certain range of humidity, corroborating similar findings in peatlands (Booth, 2002; Charman and Warner, 1997) and forest soil (Bamforth, 2015). The relationship with soil carbon content has been rarely investigated, although early works suggested that soils with contrasting carbon contents host different communities (Bonnet, 1964; Bonnet and Thomas, 1960). Therefore, Arcellinida can be considered a suitable indicator group to monitor environmental crises in the Anthropocene, linked to global change. Indeed, communities host species sensitive to modifications in climate and carbon loss in soil, widely considered as a consequence of climate change (Beillouin et al., 2023; Guo et al., 2023).

Species richness also provides information on ecosystem functioning, as functional diversity increases concomitantly (Petchey and Gaston, 2002). The final Random Forest model explained 20% of the spatial variation in OTU richness, indicating that the included environmental predictors accounted for a modest but substantial proportion of the observed patterns, while much of the variation is likely driven by unmeasured or stochastic factors. Therefore, the selected variables must be taken into account. Our Random Forest model selected drivers related to soil carbon and humidity. Organic carbon stocks (ocs) integrates multiple environmental and biological properties, such as vegetation inputs, decomposition dynamics and water retention capacity (Schmidt et al., 2011). The asymmetric, hump-shaped response of richness to organic carbon stocks (Fig. 5A) is consistent with two contrasting mechanisms acting along this gradient: the modest increase from low to intermediate stocks plausibly reflects the accumulation of prey microorganisms and microhabitats as carbon inputs build up, whereas the steep decline at high stocks likely reflects the conditions under which carbon accumulates, waterlogged, acidic and nutrient-poor, recalcitrant-rich soils such as those of Sphagnum peatlands, which most likely act as strong ecological filters on Arcellinida.

Furthermore, silt content (silt) influences soil pore architecture and water retention (Hudson, 1994). In this context, previous studies have shown that the physical architecture of the soil controls microbial diversity (Young and Crawford, 2004), as suggested in Figure 5C; and that soil moisture is one of the main factors determining the amoeba communities (Booth, 2002; ElKhouri-Vidarte et al., 2026; Mitchell et al., 2000).

Consistently, increasing precipitation seasonality (bio15) was associated with lower OTU richness (Fig. 5; B), suggesting that greater climatic variability may reduce habitat stability for moisture-dependent taxa. Projections for Bulgaria point in this direction: seasonal changes, with a significant decrease in summer rainfall and a reduction in soil moisture, which could pose challenges for agriculture and water management (Panayotov, 2024). Therefore, our results suggest that climate change, characterized by seasonal rainfall patterns, could affect the diversity of Arcellinida, reducing its species diversity.

## CONCLUSIONS

Our results show that Arcellinida diversity changes sharply depending on the ecosystem type. This suggests that most species overlap little in their realized niches. Therefore, changes in their environment such as climate change may have considerable consequences on their diversity. Climate projections for Bulgaria indicate increasing temperatures and a stronger seasonality of precipitation, leading to reduced soil moisture during the driest season. If confirmed, these projections will pose significant challenges for public health, agriculture, and water demand (Panayotov, 2024). Our results show that Arcellinida communities diversity peaks under stable conditions of humidity in temperate climates like in Bulgaria, as they do under Mediterranean climates (ElKhouri Vidarte et al., 2026). Drought has therefore an adverse effect, and global warning may cause local extinctions in Arcellinida. Future works that incorporate climate modelling and large, continental-wide sampling designs will show if Arcellinida communities should be resilient to climate change or if they will undergo the fifth extinction crisis, just like animals and plants. The consequences of the impoverishment of these keystone organisms on the functioning of natural microbiota would then need to be evaluated.

## CREDIT AUTHORSHIP CONTRIBUTION STATEMENT

**Ángel García-Bodelón:** Conceptualization, Data curation, Formal analysis, Investigation, Methodology, Project administration, Visualization, Writing – original draft, Writing – review & editing. **Milcho Todorov:** Conceptualization, Funding acquisition, Project administration, Resources, Writing – review & editing. **Nura ElKhouri-Vidarte:** Formal analysis, Writing – review & editing. **Carmen Soler-Zamora:** Formal analysis, Visualization, Writing – review & editing. **Bertrand Fournier:** Conceptualization, Methodology, Formal analysis, Supervision, Validation, Writing – review & editing. **Enrique Lara:** Conceptualization, Funding acquisition, Methodology, Project administration, Supervision, Validation, Writing – review & editing.

## DATA AVAILABILITY

All tables and figures, including the supplementary tables and figures associated with this study, are publicly available on Zenodo (https://doi.org/10.5281/zenodo.23013955).

## ACKNOWLEDGEMENTS

This work was supported by the “PIPF-2023/ECO-29443” program from the Comunidad de Madrid (Spain) to Á. García-Bodelón and also by the MYXOTROPIC VII project given to E. Lara by the Spanish Government PID2021-128499NB-I00 https://doi.org/10.13039/501100011033/, (MCIU/AEI/FEDER, UE). The authors would also like to thank Dr Jesús Muñoz for his valuable advice.

