## Supplementary material for "Precipitation seasonality and soil carbon shape Arcellinida diversity across Bulgaria’s ecosystems": Suplementary data

| Supplementary tables | |
| --- | --- |
| Figure S1 | Page 2 |
| Figure S2 | Page 2 |
| Table S1 | Page 3 |
| Table S2 | Page 4 |
| Table S3 | Page 4 |

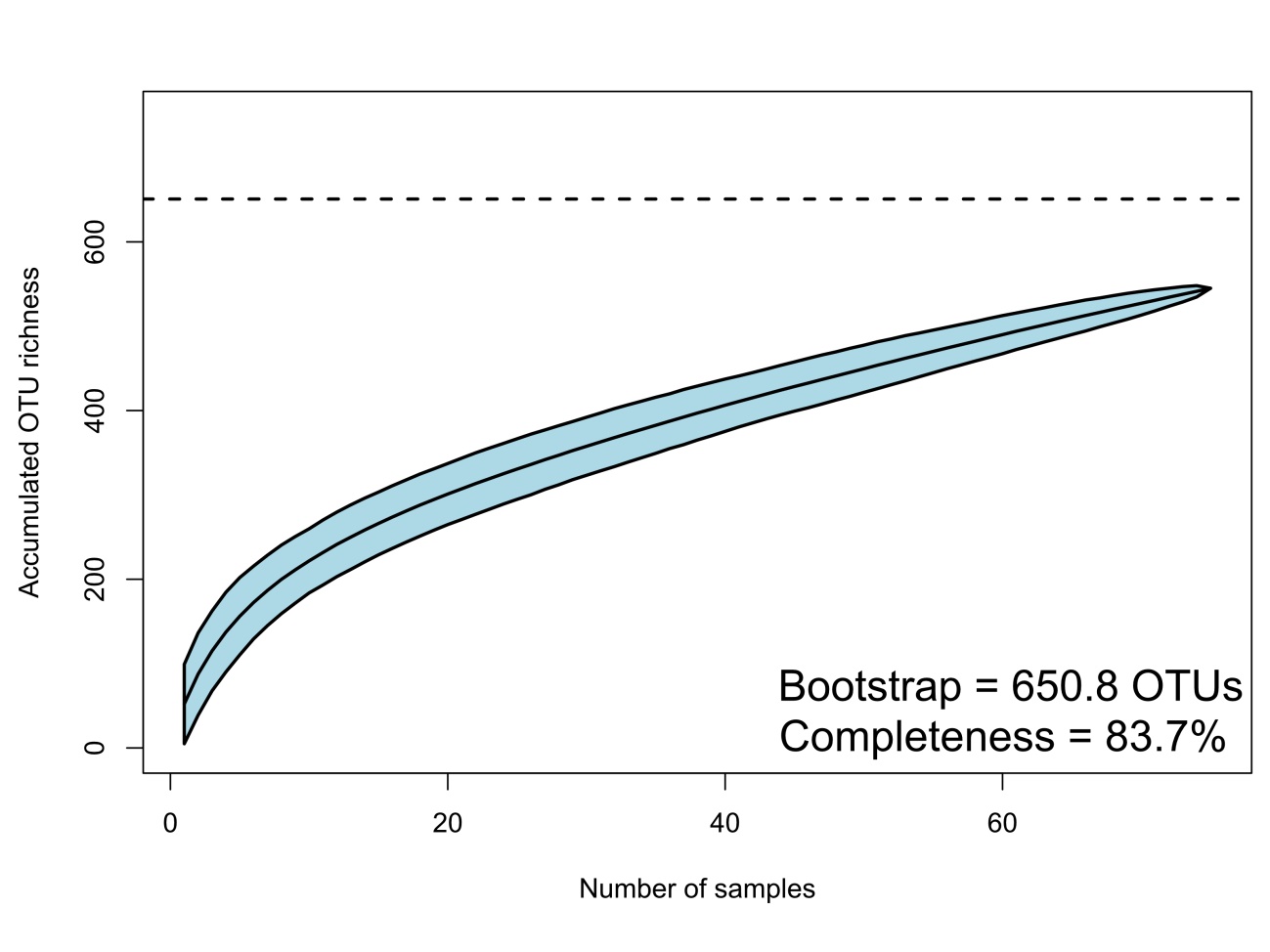

Figure S1. OTU accumulation curve across all habitats. The proportion of estimated OTU richness detected was calculated as the ratio between observed OTU richness and bootstrap-estimated richness, shown by the dotted line.

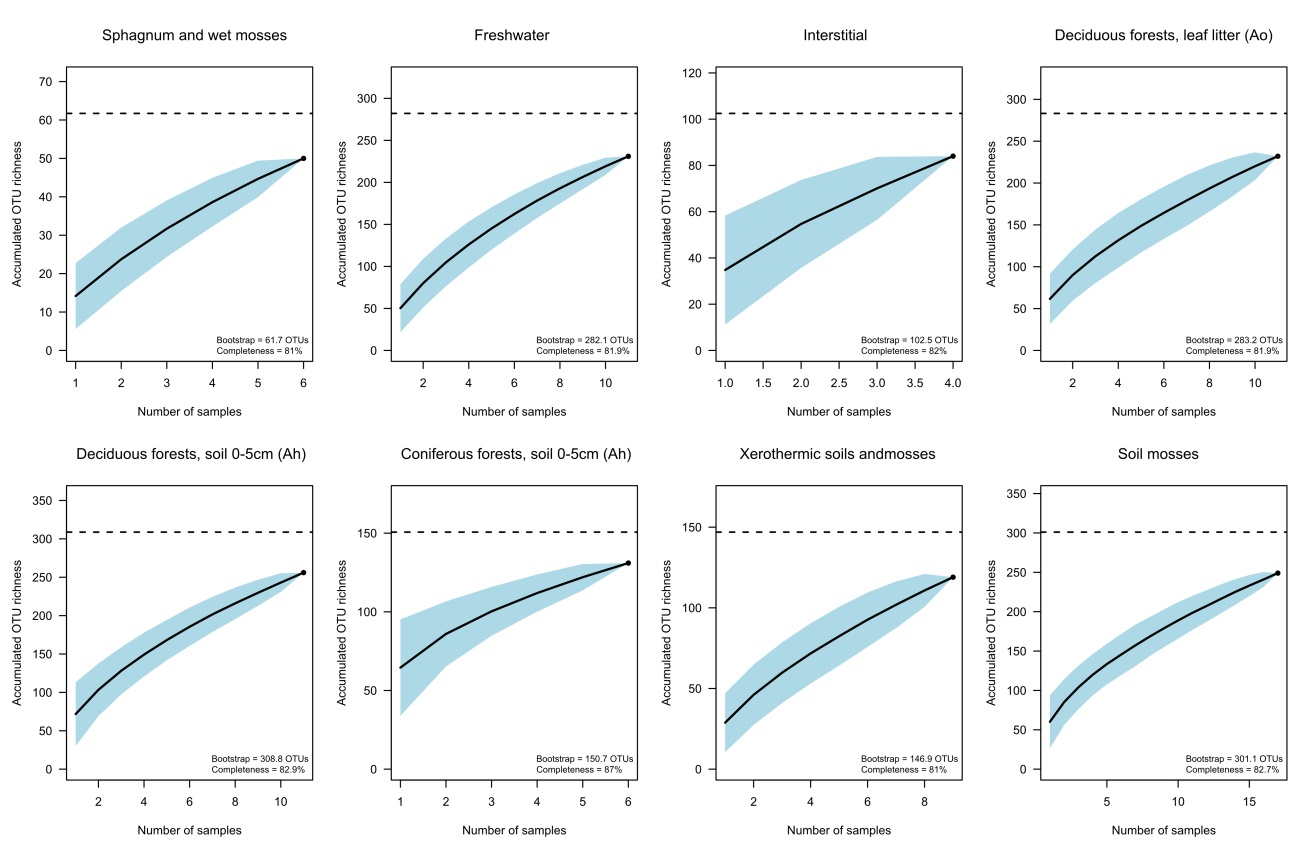

Figure S2. OTU accumulation curves by habitat. The proportion of estimated OTU richness detected was calculated as the ratio between observed OTU richness and bootstrap-estimated richness, shown by the dotted line.

Table S1. Number of Illumina reads per sample through the DADA2 pipeline.

| Sample | input | filtered | denoisedF | denoisedR | merged | nonchim | percentage |
| --- | --- | --- | --- | --- | --- | --- | --- |
| Sample 1 | 274929 | 241277 | 239916 | 239567 | 225270 | 225018 | 81.85 |
| Sample 3 | 341594 | 293539 | 291243 | 292237 | 282560 | 281947 | 82.54 |
| Sample 4 | 258029 | 223369 | 222403 | 222447 | 213538 | 212876 | 82.5 |
| Sample 5 | 282896 | 241143 | 239599 | 239832 | 235486 | 233551 | 82.56 |
| Sample 6 | 484069 | 421209 | 418392 | 419424 | 403853 | 398701 | 82.36 |
| Sample 7 | 470676 | 408663 | 405523 | 406868 | 393141 | 386575 | 82.13 |
| Sample 8 | 431331 | 366307 | 361298 | 362787 | 343731 | 324741 | 75.29 |
| Sample 9 | 380165 | 330440 | 327038 | 326280 | 298230 | 290831 | 76.5 |
| Sample 10 | 956776 | 800405 | 792794 | 793698 | 765240 | 689069 | 72.02 |
| Sample 11 | 857369 | 710822 | 708074 | 707855 | 694828 | 679192 | 79.22 |
| Sample 12 | 861561 | 765807 | 758027 | 758988 | 708553 | 667118 | 77.43 |
| Sample 13 | 452966 | 374126 | 370841 | 370484 | 354733 | 347665 | 76.75 |
| Sample 15 | 298352 | 252835 | 249318 | 248593 | 234909 | 229926 | 77.07 |
| Sample 16 | 915157 | 754208 | 748938 | 748999 | 727359 | 695734 | 76.02 |
| Sample 17 | 482656 | 309025 | 305374 | 305597 | 255728 | 242685 | 50.28 |
| Sample 18 | 485222 | 360728 | 357195 | 356859 | 325011 | 309775 | 63.84 |
| Sample 19 | 10646 | 8605 | 8172 | 8011 | 6759 | 6321 | 59.37 |
| Sample 20 | 635499 | 503085 | 500889 | 500077 | 483710 | 478084 | 75.23 |
| Sample 21 | 434866 | 360136 | 357716 | 358574 | 348347 | 337097 | 77.52 |
| Sample 22 | 721488 | 572931 | 568440 | 569378 | 530536 | 509586 | 70.63 |
| Sample 23 | 475175 | 394225 | 391981 | 391045 | 371642 | 340482 | 71.65 |
| Sample 25 | 688520 | 601690 | 596826 | 595159 | 557940 | 553040 | 80.32 |
| Sample 26 | 1004411 | 874325 | 870390 | 868269 | 836695 | 768637 | 76.53 |
| Sample27 | 1051608 | 921908 | 915927 | 915147 | 879234 | 822743 | 78.24 |
| Sample 29 | 8449629 | 7415895 | 7374529 | 7397541 | 6575622 | 5982981 | 70.81 |
| Sample 30 | 639841 | 551902 | 545489 | 546344 | 511969 | 506142 | 79.1 |
| Sample 31 | 916461 | 797380 | 791413 | 792014 | 741885 | 721788 | 78.76 |
| Sample 32 | 322630 | 287868 | 286719 | 286705 | 277491 | 275546 | 85.41 |
| Sample 33 | 516000 | 428775 | 425517 | 425131 | 390843 | 376455 | 72.96 |
| Sample 34 | 460605 | 401890 | 398508 | 398802 | 385594 | 371435 | 80.64 |
| Sample 35 | 607859 | 540912 | 535151 | 536527 | 508731 | 487958 | 80.27 |
| Sample 36 | 358816 | 315109 | 312886 | 313254 | 301329 | 295444 | 82.34 |
| Sample 37 | 783422 | 667783 | 662510 | 663371 | 635645 | 610358 | 77.91 |
| Sample 38 | 8235 | 5476 | 5127 | 5074 | 3662 | 3469 | 42.13 |
| Sample 39 | 650697 | 550420 | 548235 | 547352 | 517787 | 484637 | 74.48 |
| Sample 40 | 16935 | 13277 | 12843 | 12685 | 10308 | 10150 | 59.94 |
| Sample 41 | 849688 | 754032 | 748854 | 748921 | 720099 | 676125 | 79.57 |
| Sample 42 | 855962 | 759716 | 755284 | 755511 | 730957 | 694314 | 81.12 |
| Sample 43 | 723230 | 628022 | 624295 | 625324 | 602535 | 571415 | 79.01 |
| Sample 44 | 821635 | 704518 | 698387 | 700761 | 655595 | 627400 | 76.36 |
| Sample 45 | 577201 | 481400 | 479122 | 479003 | 458046 | 447617 | 77.55 |
| Sample 46 | 494291 | 426981 | 422240 | 423771 | 396192 | 376145 | 76.1 |
| Sample 47 | 666591 | 582703 | 579578 | 580040 | 553267 | 537223 | 80.59 |
| Sample 48 | 339653 | 276995 | 275114 | 275012 | 263826 | 258908 | 76.23 |
| Sample 49 | 274195 | 235438 | 234217 | 234300 | 224260 | 215091 | 78.44 |
| Sample 50 | 595727 | 524366 | 523032 | 522810 | 507335 | 465587 | 78.15 |
| Sample 51 | 296951 | 262618 | 261898 | 261551 | 247800 | 235746 | 79.39 |
| Sample 52 | 584555 | 515420 | 513982 | 513358 | 492780 | 452884 | 77.48 |
| Sample 53 | 585319 | 517160 | 512960 | 514100 | 491868 | 486315 | 83.09 |
| Sample 55 | 264172 | 226326 | 223406 | 224277 | 210543 | 206484 | 78.16 |
| Sample 56 | 425707 | 359907 | 357776 | 358145 | 322513 | 319166 | 74.97 |
| Sample 57 | 616667 | 538945 | 534053 | 536046 | 497798 | 486058 | 78.82 |
| Sample 58 | 135790 | 113898 | 112967 | 113010 | 109032 | 107691 | 79.31 |
| Sample 59 | 321272 | 277854 | 276368 | 276687 | 267908 | 258424 | 80.44 |
| Sample 60 | 335671 | 264919 | 263224 | 262195 | 160770 | 154295 | 45.97 |
| Sample 62 | 657130 | 583571 | 580683 | 580427 | 553344 | 515006 | 78.37 |
| Sample 63 | 739714 | 633322 | 630977 | 630358 | 585361 | 571608 | 77.27 |
| Sample 64 | 644818 | 474873 | 471652 | 470524 | 379604 | 352149 | 54.61 |
| Sample 65 | 709022 | 620390 | 618732 | 616986 | 605881 | 561346 | 79.17 |
| Sample 66 | 798319 | 704088 | 699809 | 701095 | 684749 | 597295 | 74.82 |
| Sample 67 | 831808 | 739511 | 733757 | 736180 | 705443 | 684194 | 82.25 |
| Sample 68 | 763610 | 682314 | 675229 | 678043 | 644985 | 556690 | 72.9 |
| Sample 69 | 723435 | 642693 | 637092 | 639127 | 598765 | 586730 | 81.1 |
| Sample 70 | 837154 | 741841 | 737196 | 736830 | 686550 | 657840 | 78.58 |
| Sample 71 | 723119 | 635634 | 631143 | 629695 | 594120 | 553667 | 76.57 |
| Sample 72 | 796241 | 690994 | 688383 | 688235 | 660377 | 596580 | 74.92 |
| Sample 73 | 685571 | 599606 | 590162 | 593808 | 551915 | 519524 | 75.78 |
| Sample 74 | 685616 | 612993 | 610424 | 610288 | 595310 | 525691 | 76.67 |
| Sample 75 | 718129 | 644423 | 639643 | 641620 | 621414 | 587817 | 81.85 |
| Sample 76 | 670112 | 587865 | 586175 | 585533 | 569637 | 540711 | 80.69 |
| Sample 77 | 650255 | 570223 | 566369 | 566044 | 536999 | 519859 | 79.95 |
| Sample 78 | 587476 | 518040 | 515839 | 514720 | 488997 | 470910 | 80.16 |
| Sample 79 | 275489 | 237992 | 236817 | 236602 | 215097 | 203210 | 73.76 |
| Sample 80 | 141226 | 123947 | 123315 | 123176 | 119051 | 113522 | 80.38 |
| Sample 81 | 390138 | 348658 | 346819 | 347304 | 340459 | 307584 | 78.84 |

Table S2. This table is provided as a separate file in CSV format (Table S2.csv)

Table S3. List of the 34 bioclimatic variables obtained from CHELSA v2.1 and the 11 edaphic variables from ISRIC databases, including their full names, corresponding abbreviations and conventional units.

|  | Variable names | Abbreviations | Conventional units |
| --- | --- | --- | --- |
| Bioclimatic variables | Mean Annual Near-Surface Air Temperature | bio1 | °C |
|  | Mean Diurnal Near-Surface Air Temperature Range | bio2 | °C |
|  | Isothermality | bio3 | °C |
|  | Temperature Seasonality | bio4 | °C/100 |
|  | Mean Daily Maximum Near-Surface Air Temperature of the Warmest Month | bio5 | °C |
|  | Mean Daily Minimum Near-Surface Air Temperature of the Coldest Month | bio6 | °C |
|  | Annual Daily Mean Near-Surface Air Temperature Range | bio7 | °C |
|  | Mean Daily Near-Surface Air Temperature of the Wettest Quarter | bio8 | °C |
|  | Mean Daily Near-Surface Air Temperature of the Driest Quarter | bio9 | °C |
|  | Mean Daily Mean Near-Surface Air Temperature of the Warmest Quarter | bio10 | °C |
|  | Mean Daily Mean Near-Surface Air Temperature of the Coldest Quarter | bio11 | °C |
|  | Annual Precipitation | bio12 | kg m-2 year-1 |
|  | Precipitation of the Wettest Month | bio13 | kg m-2 month-1 |
|  | Precipitation of the Driest Month | bio14 | kg m-2 month-1 |
|  | Precipitation Seasonality | bio15 | kg m-2 |
|  | Mean Monthly Precipitation of the Wettest Quarter | bio16 | kg m-2 month-1 |
|  | Mean Monthly Precipitation of the Driest Quarter | bio17 | kg m-2 month-1 |
|  | Mean Monthly Precipitation of the Warmest Quarter | bio18 | kg m-2 month-1 |
|  | Mean Monthly Precipitation of the Coldest Quarter | bio19 | kg m-2 month-1 |
|  | Monthly Climate Moisture Index | cmi | kg m-2 month-1 |
|  | Growing Degree Days Heat Sum above 0 °C | gdd0 | °C |
|  | First Day of the Growing Season TREELIM | fgd | julian day |
|  | Growing Season Length | gsl | days |
|  | Accumulated Precipiation Amount on Growing Season Days | gsp | kg m-2 gsl-1 |
|  | Mean Temperature of Growing Season Days | gst | °C |
|  | Near-Surface Relative Humidity | hurs | % |
|  | Number of Growing Degree Days above 0 °C | ngd0 | number of days |
|  | Net Primary Production on Land as Carbon Mass Flux | npp | g C m-2 yr-1 |
|  | Monthly Potential Evapotranspiration | pet | kg m-2 month-1 |
|  | Surface Downwelling Shortwave Flux in Air | rsds | MJ m-2 |
|  | Snow Cover days | scd | days |
|  | Near-Surface Wind Speed | sfcWind | m s-1 |
|  | Soil Water Balance | swb | kg m−2 yr−1 |
|  | Vapor Pressure Deficit | vpd | Pa |
| Edaphic variables | Bulk density | bdod | kg/dm3 |
|  | CEC buffered at pH7 | cec | cmol(c)/kg |
|  | Coarse fragments | cfvo | cm3/100cm3 (vol%) |
|  | Clay | clay | g/100g (%) |
|  | Nitrogen | nitrogen | g/kg |
|  | Organic carbon density | ocd | kg/m3 |
|  | Organic carbon stocks | ocs | kg/m2 |
|  | Soil organic carbon | soc | g/kg |
|  | pH water | phh2o | - |
|  | Sand | sand | g/100g (%) |
|  | Silt | silt | g/100g (%) |
